# Neuromuscular fatigue reduces responsiveness when controlling leg external forces

**DOI:** 10.1101/2023.05.24.541485

**Authors:** Pawel Kudzia, James M. Wakeling, Stephen N. Robinovitch, J. Maxwell Donelan

**Author notes:** **Corresponding author:** J. Maxwell Donelan, Department of Biomedical Physiology and Kinesiology, Shrum Science Centre K9625, 8888 University Drive, Burnaby, BC V5A 1S6, Canada.

## Abstract

In legged movement, our legs push against the ground, generating external force vectors that enable agile movements. Neuromuscular fatigue can reduce agility by causing physiological changes, such as slowing muscle reaction time, altering proprioception, and delaying neuromuscular control. Fatigue may deteriorate the nervous system’s control of leg external forces, contributing to reductions in agility. In this study, we investigated the effect of fatigue on the performance of the nervous system in controlling the vertical component of leg external force ground reaction forces. We hypothesized that increased leg fatigue would lead to declines in both the responsiveness (speed) and accuracy of leg force control. To test this hypothesis, we used an apparatus that allowed participants to exert controlled vertical forces with one leg against a force plate while immobilizing the rest of their bodies. Participants adjusted their leg external force to match step targets displayed on a screen. We induced fatigue by having participants maintain submaximal leg forces, and we measured leg force control performance between fatigue trials. Results showed a significant 26% reduction in mean maximum force production, leading to a substantial decline in leg force control responsiveness, as evidenced by a 23% increase in rise time and a 25% narrowing of bandwidth. However, fatigue did not significantly reduce leg force control accuracy. Understanding the effects of fatigue on leg force control can inform the development of strategies and technologies to sustain agile performance, even in the presence of fatigue.

**New and Noteworthy:** We developed a method to probe the influence of neuromuscular fatigue on the control of leg external forces. Our findings demonstrate that while fatigue significantly diminishes responsiveness (speed), it does not compromise the accuracy of control. These insights enhance our understanding of legged agility and could guide the development of strategies for optimizing leg force control performance. This study paves the way for future research aimed at identifying and employing effective strategies to maintain agility in the face of fatigue.

## Introduction

One way that legged animals, including humans, demonstrate agility is through effective control of external leg forces. We can push against the ground with any part of our bodies, but we often do this using our legs to push our feet against the ground. The resulting reaction force from the environment, a vector quantity, accelerates our bodies. Here we consider the rapid and accurate control of the external force vector to be agility [1]. Examples of legged agility include accelerating from a standstill, jumping high and far over obstacles, and quickly changing movement direction, all of which require effective control of our leg’s external forces.

Neuromuscular fatigue reduces agility. Fatigue leads to a decline in muscle force-generating capabilities [2] and reductions in maximum muscle shortening velocities [3], [4]. As mechanical power is the product of both force and shortening velocity, fatigue reduces maximum muscle power [2]. The effect of fatigue on muscle power and agility is well-documented in sports performance, particularly in rapid change-of-direction sports like soccer and basketball [5]–[7]. For example, fatigue reduces power in jumping leading to jumps of shorter distances and lower heights [8]–[12]. A second example is sprinting, in which high performance relies on fast-twitch muscle fibres. These same fibres are also the most susceptible to fatigue, leading to declines in speed in the final meters of a 100-m race [13]–[16]. Other physiological changes that may arise from fatigue include slowing of muscle reaction time [17], altering of proprioception [18], and delaying our neuromuscular control in ways that reduce the control of muscles [19]. The physiological changes described are some of the mechanisms that may explain reductions in agility and deteriorate the nervous system’s control of the leg’s external forces. In support of this idea, soccer players show a reduced ability to complete legged balancing tasks on unstable surfaces when fatigued [20].

The aim of our study was to characterize the effect of fatigue on the nervous system’s ability to rapidly and accurately control vertical external leg forces. We hypothesized that increased leg fatigue would deteriorate the leg’s control of submaximal vertical external forces, as characterized by both reduced responsiveness and accuracy. We refer to responsiveness as to how quickly a person can control the magnitude of leg vertical external force when presented with a target force level while we refer to accuracy as how close and variable their controlled force is to and around an intended target value. To test this hypothesis, we built a custom apparatus that constrained participants’ bodies from moving, allowing their legs to maintain a static posture while controlling the external vertical force magnitude below their feet. We fatigued participants by having them hold submaximal vertical leg forces equal to 25% of their maximum voluntary external force and measured the leg’s force control performance between fatigue trials in matching commanded changes in force. We quantified fatigue as reductions in vertical force during maximal voluntary leg contractions, increases in force variability, and a shift to lower mean frequencies in leg muscle activity. To quantify force control performance, we instructed participants to best match visually displayed target step changes in vertical force magnitude by pushing more or less against a ground-mounted force plate. We evaluated control performance based on how responsive and accurate participants were in matching their leg’s external force vertical magnitude to the commanded target changes in vertical force.

## Methods

### Participants

We recruited 18 participants for the study (identifying as female: n = 11, identifying as male =7; age: 26.5±3.8y; body mass:74±16 kg; height: 171±12 cm; shoe size: 9+2 US sizing; mean±std). The Office of Research Ethics at Simon Fraser University approved the study. Participants provided us with verbal and written informed consent before participating.

### Experimental Design

To characterize human leg force control, we tested the step response of participants as they selectively controlled external leg force. We used a custom apparatus (Figure 1) that we had previously built [1]. The apparatus consisted of a ground-embedded force plate (Bertec Corporation, Ohio, USA) that participants stood on, constrained in both the vertical and horizontal directions. The force plate was connected to a computer through a data acquisition unit (USB-6229, National Instruments Corporation, Texas, USA), which sampled data at 1000 Hz. Participants were able to exert a variable but controlled external force vector onto the ground by selectively pushing down with their leg. Our device has rigid supports that constrain participants at the shoulders and forearms preventing movement vertical and horizontal motion of their upper body. Although control of force-position in the medial-lateral and anterior-posterior directions (i.e., center of pressure control) and control of all three orthogonal force magnitude components of the external force vector is important, our work here focused only on the control of vertical force magnitude. In our prior work, we characterized the control of different step sizes of medial-lateral and anterior-posterior force positions, as well as the control of a range of sub-maximal vertical force magnitudes and found negligible differences in control characteristics for these different components of the external force vector [1]. By focusing on only the vertical component of force, we believe our findings will represent the range of control characteristics for all aspects of controlling the external force vector at submaximal forces and any control changes resulting from fatigue.

**Figure 1:**
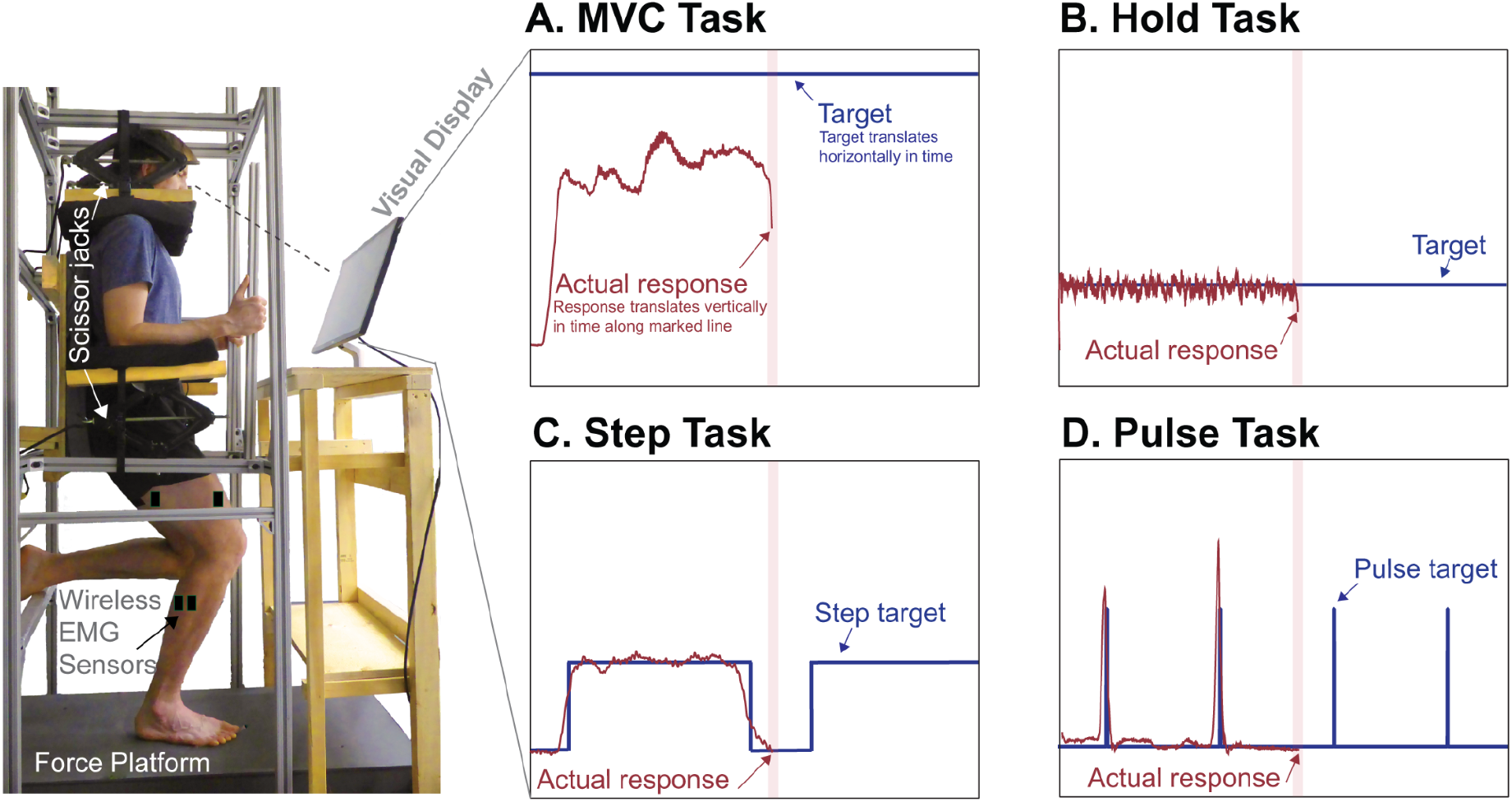
Apparatus used to characterize external force control and fatigue participants. The setup consisted of adjustable scissor jacks that constrained the vertical and horizontal motion of the participants’ torso, pressing down on their shoulders and against their forearms, and a stiff 80/20 aluminum frame securely mounted to the ground. Each participant stood on their right leg in a posture resembling the mid-stance phase of a run and applied a vertical force to the ground by pushing down with their foot onto a force platform. The real-time feedback displayed to the participants showed the vertical force they were applying to the ground, as well as the target they were trying to match. There were four different tasks with four distinct visual targets: A. the MVC Task, which involved pushing down maximally for 10 seconds while trying to reach a motivating but unattainable target (5.0 BW); B. the Hold Task, which required participants to match a target equivalent to 1.4 times their body weight (1.4 BW) for 73 seconds; C. the Step Task, which involved rapidly and accurately controlling vertical force to match 10 upcoming step targets in 73 seconds; and D. the Pulse Task, which required participants to rapidly match 10 upcoming step targets in 73 seconds by changing their vertical force to match a target force that approximated a pulse. Participant shown in figure is author and has given consent for his picture to be used in this publication.

To objectively quantify fatigue, we recorded electromyography (EMG) from the leg muscles of each participant. We placed wireless EMG sensors (Delsys, Natick, USA) on the middle of the muscle belly on the vastus medialis, vastus lateralis, rectus femoris, bicep femoris, gastrocnemius medialis, and gastrocnemius lateralis of the right leg muscles of each participant. Prior to placing the sensors, we prepared the location by removing any excess hair using a razor and rubbing it down with an alcohol swab, following standard procedures. We synchronized our EMG system with our force plate using a custom code, which triggered the collection of both force plate and EMG data. We used an EMG sampling rate of 2000 Hz.

After placing the EMG sensors, we fitted participants into the experimental apparatus. When fitting participants, we adjusted the apparatus such that participants maintained an approximately 15-degree knee flexion angle in their right leg, a posture similar to that assumed by a runner at the start of the stance phase [1], [21]. In this fitting process, we moved components of the apparatus to push down on the participant’s shoulders and up on their arms by manually rotating four scissor jacks (Figure 1). Our goal here was to ensure that participants were fixed tightly within the apparatus and unable to move side-to-side or vertically.

To assess the effects of leg fatigue on the control of leg external force, we commanded four types of force control tasks. In all tasks, participants stood on their right leg, unless otherwise instructed. We used visual feedback to allow participants to compare their actual vertical force magnitude to a target that we instructed them to match. We provided real-time visual feedback using a monitor, where our custom-developed software displayed both the real-time feedback of the vertical force magnitude signal that the participant’s foot exerted onto the force plate and the target that the participant tried to rapidly and accurately match. For visual display, we normalized the real-time signal to each participant’s body weight and filtered the raw force signal using a low-pass fourth-order Butterworth filter with a cut-off frequency of 10 Hz. We programmed the real-time signal to display at the center of the screen and constrained it to only move up and down as the participant pushed more or less against the force plate. We programmed the target to slide past the real-time signal, giving participants time to view any upcoming changes in the target before they occurred.

In the first force control task, referred to as the Maximum Voluntary Contraction Task (MVC Task), we evaluated the maximum effort of voluntary force that the leg could exert on the ground. To collect the MVC Task, we asked participants to push down against the ground as hard as possible and try to reach an unattainable target of 5 times their body weight (5.0 BW) displayed on the screen. From pilot experiments, we found that providing participants with a target to reach, even if it was not attainable, motivated them to push harder to try and reach it. Each MVC Task lasted 10 seconds as participants pushed down with their foot against the ground as hard as they could.

We designed the second force control task, referred to as the Hold Task, to fatigue each participant’s right leg. In the Hold Task, we asked participants to match and hold a target equal to their body weight plus 25% of their pre-fatigue maximum voluntary contraction, as determined in the MVC Task before the onset of any leg fatigue [22]. This type of sustained voluntary contraction progressively recruits more motor units and leads to muscle fatigue in the active muscles [22], [23]. For each Hold Task, we asked participants to hold their force level to the best of their ability to match the target for a total duration of 73 seconds, the same duration as the third leg force control task.

In the third leg force control task, referred to as the Step Task, we probed force control characteristics by commanding target step changes in leg vertical force magnitude. Because both initial speed to the new target and steady-state accuracy around the new target are objectives, we anticipate that the nervous system will depend on feedback control to accomplish this task. In a single Step Task, the target step function consisted of a square wave of ten matching size target steps, each 4 seconds in duration, with 3 seconds between steps (totaling 73 seconds per task). The lower value of the step target was 1.0 BW and the upper value (i.e., the size of the target) was 1.4 BW. We selected the target step size based on a previous experiment where we studied step responses by commanding a range of target step sizes, both smaller and larger than the magnitude chosen here [1]. We observed minor changes in control characteristics between targets of varying step sizes, so we chose an intermediate step size for this experiment.

In the fourth force control task, which we referred to as the Pulse Task, we aimed to probe rapid force control by asking participants to respond as rapidly as possible to changes in the target. Because this task emphasizes the initial speed of response over steady-state accuracy, we anticipate that the nervous system will emphasize feedforward control over feedback control when accomplishing this task [24]. The target in this task resembled a pulse, changing momentarily for 0.05s from 1.0 BW to the target step size of 1.4 BW (the same step size as in the Step Task). In a single Pulse Task, we programmed the target pulse to go from body weight to the target step size ten times over the duration of 73 seconds.

## Experimental Protocol

We conducted the experiment in a single session. The session included five conditions: Training, Pre-Fatigue, Fatigue-1, Fatigue-2, and Fatigue-3 (Figure 2). We started the study with the Training condition to familiarize participants with the experiment. In this condition, we explained and demonstrated how the visual feedback and force plate pushing worked. Then, we asked each participant to perform two repetitions of the MVC Task followed by three repetitions of the Step Task. From our previous work, we believe this is enough training in the apparatus to eliminate any learning effects. After the Training condition, we gave participants a 1-minute break to stand freely on both feet in the apparatus. Next, in the Pre-Fatigue condition, we had participants perform two repetitions of the MVC Task followed by a 1-minute break, then three Step Tasks followed by a 1-minute break, and then two Pulse Tasks. We then gave participants a 1-minute break to rest before the next condition. Next, we started the sequence of three fatigue conditions. Each Fatigue condition consisted of a Hold Task followed by an MVC Task, which was repeated three times within each Fatigue condition. After the third and final MVC Task in each Fatigue condition, we immediately had each participant perform three repetitions of the Step Task to probe force control. For each Fatigue condition, we repeated this same sequence of tasks. On the third Fatigue condition and after the final Step Task, participants performed two Pulse Tasks. We gave participants a 1-minute break between Fatigue conditions.

**Figure 2:**
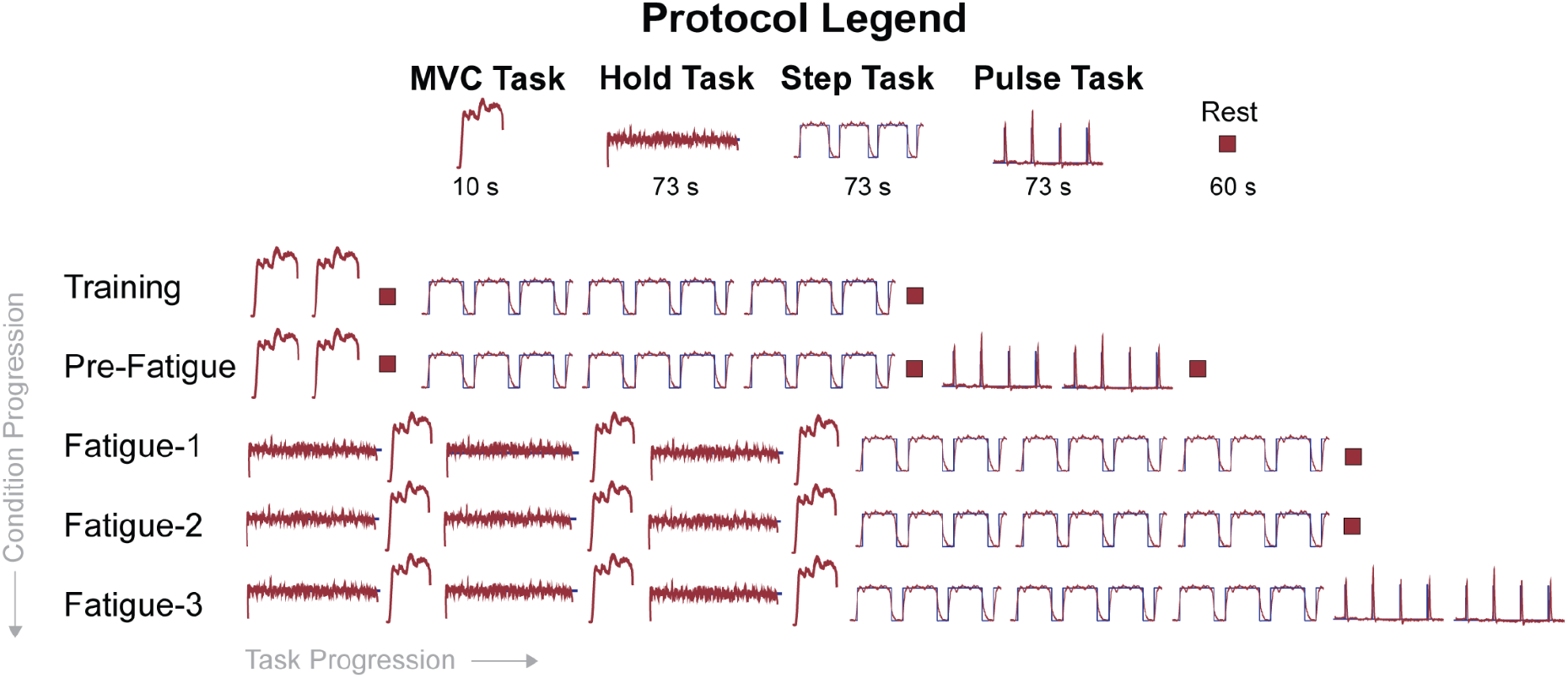
A schematic of the experimental protocol which consisted of 5 conditions: Training, Pre-Fatigue, First Fatigue, Middle Fatigue, and Final Fatigue Each condition was composed of a combination of our force control tasks: Maximum Voluntary Contraction MVC Task, Hold Task, Step Task, and Pulse Task. The progression within each condition is shown as going from left to right and the progression of conditions goes from top to bottom starting from Training and finishing with Final Fatigue.

### Data Analysis

For each MVC Task, we evaluated the mean vertical force output as one objective measure of fatigue (Figure 3-A). We quantified the mean vertical force output by taking the average of the signal during the midway point of the MVC Task between the 4-8 second time interval.

**Figure 3:**
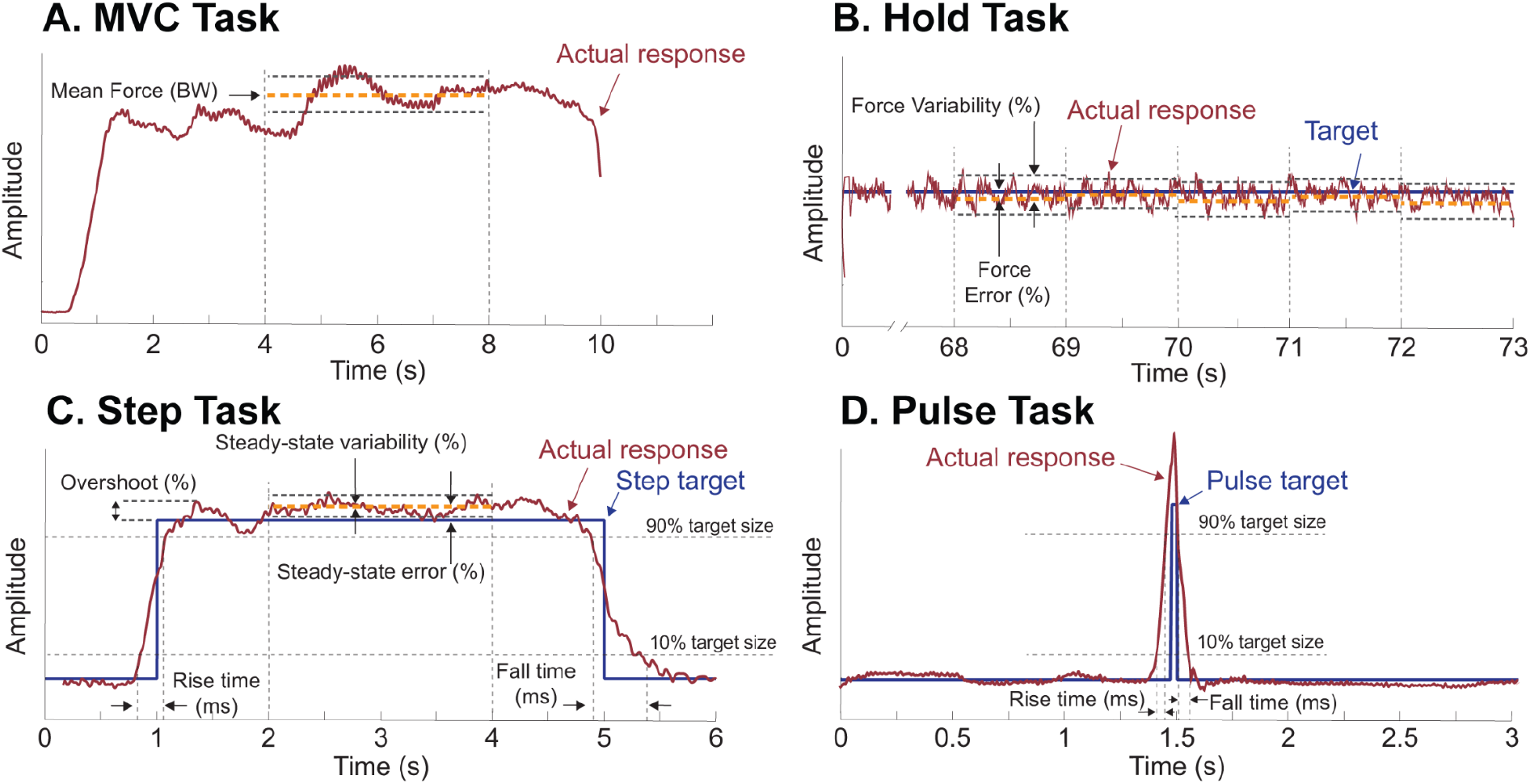
A visual analysis of the four leg force control tasks evaluated during the experiment. A. In the MVC Task, we evaluated the mean force of the signal between 4-8 seconds. B. In the Hold Task, we omitted any analysis during the first 5 seconds but calculated force variability for each 1-second interval for the duration of the task. C. We evaluated the responsiveness (rise time, fall time, bandwidth) and accuracy (overshoot, steady-state error, steady-state variability) for each step response in the Step Task. D. For each Pulse target in the Pulse Task we evaluated the responsiveness.

For each Hold Task, we evaluated the force variability of the response (Figure 3-B). We segmented each Hold Task data into one-second segments to quantity changes over time. We omitted any further analysis from the first 5 seconds of this signal as participants adjusted their vertical force to match the target during this time frame. For each 1-second segment, we then quantified force variability by calculating the standard deviation of the signal during each 1-second segment, dividing this by the mean of the signal during that time frame and multiplying this value by 100. This expresses variability as a percentage of the force magnitude.

For each step response, within each Step Task, we evaluated the responsiveness in controlling leg external force (Figure 3-C). To accomplish this, we took step data from each Step Task and segmented it into ten individual step responses such that each segmentation consisted of a six-second single-step response, with one second before the step-up and one second after the step-down. For each step response, we evaluated the rise time (in units of milliseconds) as the time for the signal to go from 10% to 90% of the target step size value [1]. We calculated bandwidth (in units of Hz) by dividing 0.35 by the rise time [25]. The bandwidth measures the maximum frequency at which a target signal can change and still be accurately tracked by the controller. We remind readers that as we calculate bandwidth using rise time, bandwidth is not an independent measure of responsiveness but rather depends on any changes in rise time. We restricted our evaluation of the rise time and bandwidth of the response to the time interval of 0.5 seconds before the step target stepped up to its high value and up to one second after the step-up occurred. Finally, we quantified fall time (in units of milliseconds) as the time for the signal to go from 90% to 10% of the target step size value. We evaluated the fall time between four to six seconds when the step signal stepped down from the target step size to 1 BW. In this paper, we use the term responsiveness to refer to rise time, bandwidth, and fall time. In robotic systems, and how we think of responsiveness in this paper, a responsive system that is capable of high-fidelity tracking of a target possesses a short rise time, a wide bandwidth, and a short fall time [26].

For each step response, within each Step Task, we evaluated the accuracy of controlling leg external force (Figure 3-C). We quantified overshoot by taking the peak value reached by the signal, subtracting the value of the target step, dividing this by the size of the step, and multiplying by 100. We expressed overshoot as the percentage that the signal exceeds the target step size value. A large overshoot can indicate under-damping in the controller—it can take longer for the system to settle into a steady state, but it may be faster to reach its target step size [1]. We restricted our evaluation of the overshoot of the response to the time interval of 0.5 seconds before the step target stepped up and to one second after the step-up occurred. Next, we quantified steady-state error by calculating the mean value of the signal between 2-4 seconds, subtracting the value of the target step size, and taking the absolute value. We divided this by the value of the target step size and multiplied it by 100 to express the steady-state error as a percentage of the target value. The steady-state error informs us how much error the control system has once it has reached the target size value and settled in its new state. A large error can be indicative of poor control as the system is not able to well match the target value. We quantified steady-state variability by calculating the standard deviation of the signal between 2-4 seconds, dividing this by the steady-state mean and multiplying this value by 100. The steady-state variability, expressed as a percentage of the steady-state mean, informs us how variable the system is once it has reached and settled on its new state. From pilot studies, we found that using the 2-4 second interval to measure steady-state error and steady-state variability was a sound assumption as participants had at this point reached a steady state. In this paper, we use the term accuracy to refer to overshoot, steady-state error, and steady-state variability—an accurate system has small overshoot, small steady-state error, and small steady-state variability. We collectively refer to the rise time, bandwidth, overshoot, steady-state error, steady-state variability, and fall time as the step response characteristics.

For each Pulse Task, we evaluated the responsiveness of the response (Figure 3-D). To accomplish this, we took the data from each Pulse Task and segmented it into ten individual step responses such that each segmentation consisted of a 3-second single-step response, with 1.5 seconds before the step-up and 1.5 seconds after the step-down. Following the same definitions of responsiveness as for the Step Task, we quantified the rise time, bandwidth, and fall time for each step in every Pulse Task.

We analyzed the EMG data to quantify changes in muscle frequency, one objective measure of fatigue. We first bandpass filtered the raw EMG signals from 30 to 400 Hz using a fourth-order zero-lag Butterworth filter and bandstop filtered the signal from 57–63 Hz to remove any residual 60 Hz noise [29]. We calculated the EMG median frequencies, a common measure of local muscle fatigue [27]. To do this, we transformed the signal from the time domain into the frequency domain by employing a fast-Fourier analysis to calculate the power spectrum. Next, we calculated median power [28], [29]. A shift to lower median frequencies over time is viewed as evidence of local muscle fatigue [28], [30].

For the Step Tasks and Pulse Task, we used exclusion criteria to remove responses that may poorly describe the measured response. For the Step Task, we used the following criteria: (1) the response steps up 0.5s before the target step function steps up, (2) the response does not step up within 0.5s after the target step function steps up, (3) the response is already greater than 10% of the target step size value during the 0.5s leading up to the step-up (4) the response steps down 1s or more before the target step function steps down. For the Pulse Task, we used the following criteria: (1) the response rises up 0.25s before the pulse target steps up, and (2) the response does not fall down until greater than 0.25s after the pulse target steps down. We removed the step responses that meet these exclusion criteria from all subsequent analyses.

### Statistical Analysis

To determine the effects of our protocol on fatigue, we evaluated several objective measures. The first measure we evaluated was changes in mean force magnitude during the maximum voluntary contraction (MVC) task. A decrease in maximum vertical force is one indicator of fatigue [23]. To calculate this measure, we compared the mean force values from the final MVC tasks completed in each condition for each participant. Our second measure was force variability during each of the consecutive Hold Tasks in each fatigue condition. Increased force variability as fatigue progressed is an indicator of local muscle fatigue [31], [32]. To calculate this measure, we compared the mean force variability for the final ten seconds of each Hold Task repeated in each fatigue condition for each participant. To estimate force variability before the onset of fatigue (referred to as “Pre-Fatigue”), we also calculated the mean force variability for the first ten seconds of the Hold Task in the first Fatigue-1 condition. Our third measure was reductions in median

EMG frequency. As with force variability, we compared the mean of the median EMG frequencies determined for the first ten seconds before the onset of fatigue and the final ten seconds of each fatigue condition. To evaluate the effects of our protocol on leg external force control, we next determined step response characteristics. For all conditions, we took the mean value of each step response characteristic for the first five Step Task responses for each participant immediately following the final MVC task. We used the same approach to characterize step response characteristics for the Step Task and the Pulse Task. To determine if there were differences in mean values between conditions, we performed a repeated-measures analysis of variance. If a significant difference was found, we conducted post hoc pairwise comparisons between the group means to identify which conditions produced similar mean values. We adjusted p-values for multiple comparisons using Bonferroni corrections. In all cases, we used MATLAB’s statistical analysis toolbox and accepted p<0.05 as statistically significant. Unless otherwise stated we present all data in the text as mean± standard deviation

## Results

### Participants fatigued during the protocol

Objective measures of fatigue demonstrated that participants were fatigued during the protocol. Task data for a representative participant are shown in Figure 4. Fatigue was demonstrated by a reduction in the mean maximum MVC force (Figure 4A). As participants progressed through the protocol conditions, they exhibited significant reductions (p=1.3e-12) in the mean force magnitude during the MVC Tasks (Figure 5-A). During Pre-Fatigue, participants on average exerted a vertical mean leg force of 2.95±0.58 BW and by the final fatigue condition, participants exerted a mean force of 2.20±0.47 BW, a significant (p=4.7e-6) reduction of 26% in mean maximum force production. Next, we found significant increases in force variability (p=0.002) as participants progressed through Fatigue conditions executing the Hold Tasks (Figure 5-B). During Pre-Fatigue, participants had a mean force variability of 0.61± 0.31% and by the final condition, participants had a mean force variability of 2.83±2.93% a significant (p=0.027) increase of 368%. Finally, we found significant reductions in the mean median frequency for several muscles specifically the gastrocnemius lateralis (p=0.002), gastrocnemius medialis (p=0.0019), and the bicep femoris (p=0.003) (Figure 5-C). Comparing the Pre-Fatigue to the final fatigue condition, the gastrocnemius lateralis medialis exhibited a significant (p=0.016) reduction of 10% in frequency reducing from 126±24 Hz to 113±27 Hz; the gastrocnemius medialis exhibited a significant (p= 0.048) reduction of 10% reducing in frequency from 133±28Hz to 119±34Hz; and the bicep femoris exhibited a reduction of 11% reducing in frequency from 90±12Hz to 79±13Hz but this only approached but did not reach significance between these two conditions (p=0.074). The other three muscles (vastus lateralis, vastus medialis, and rectus) did not show any significant changes in EMG median frequency.

**Figure 4:**
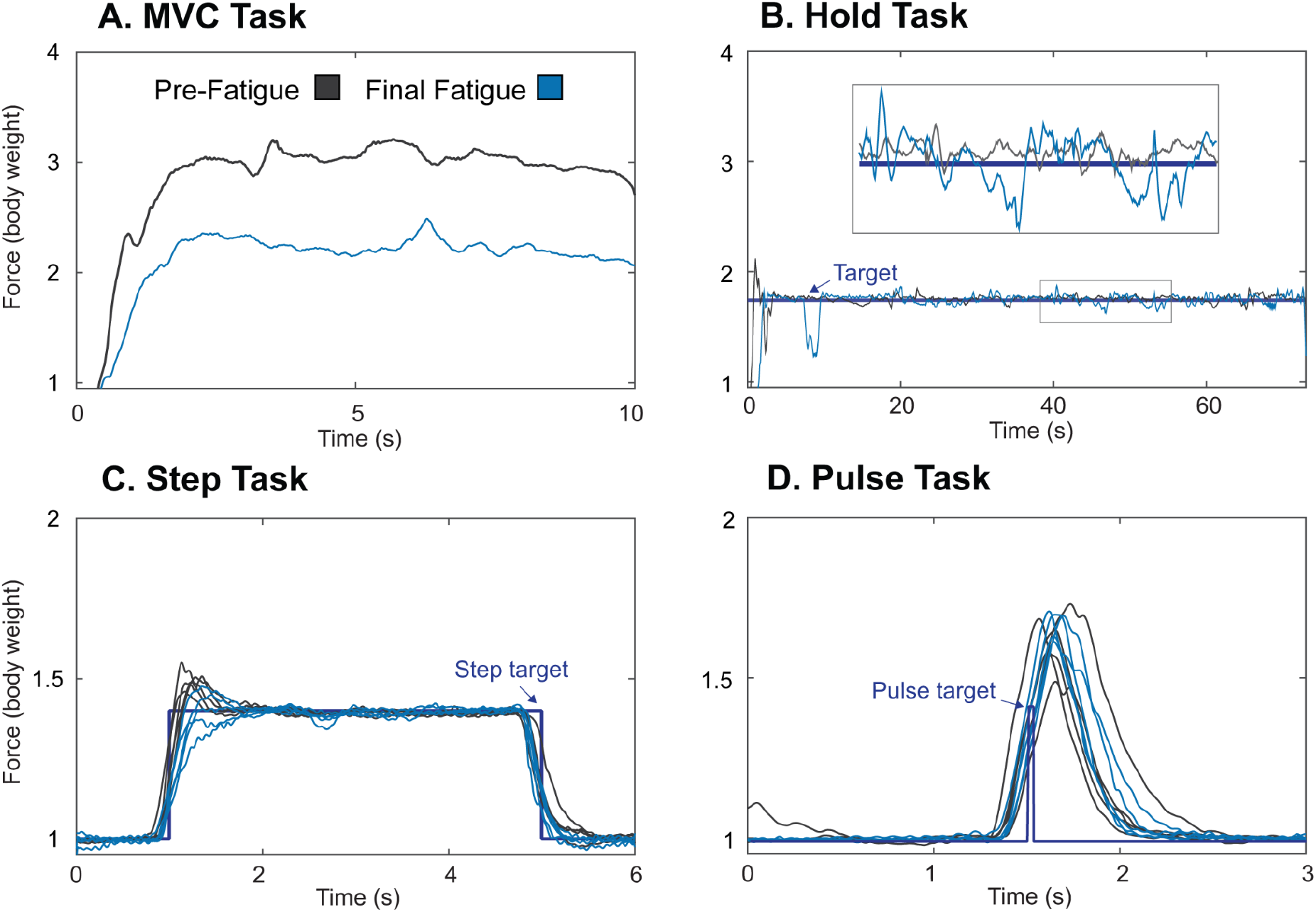
Representative data for one participant comparing the Pre-Fatigue (grey) and Final Fatigue (blue) conditions for the A. MVC Task, B. Hold Task, C. Step Task, and D. Pulse Task. For C. and D. the first five responses immediately following the final MVC Task in each condition are shown. As the participant progressed through the protocol, they exhibited fatigue as indicated by a reduction in force for the MVC task and an increase in force variability during the Hold Task.

**Figure 5:**
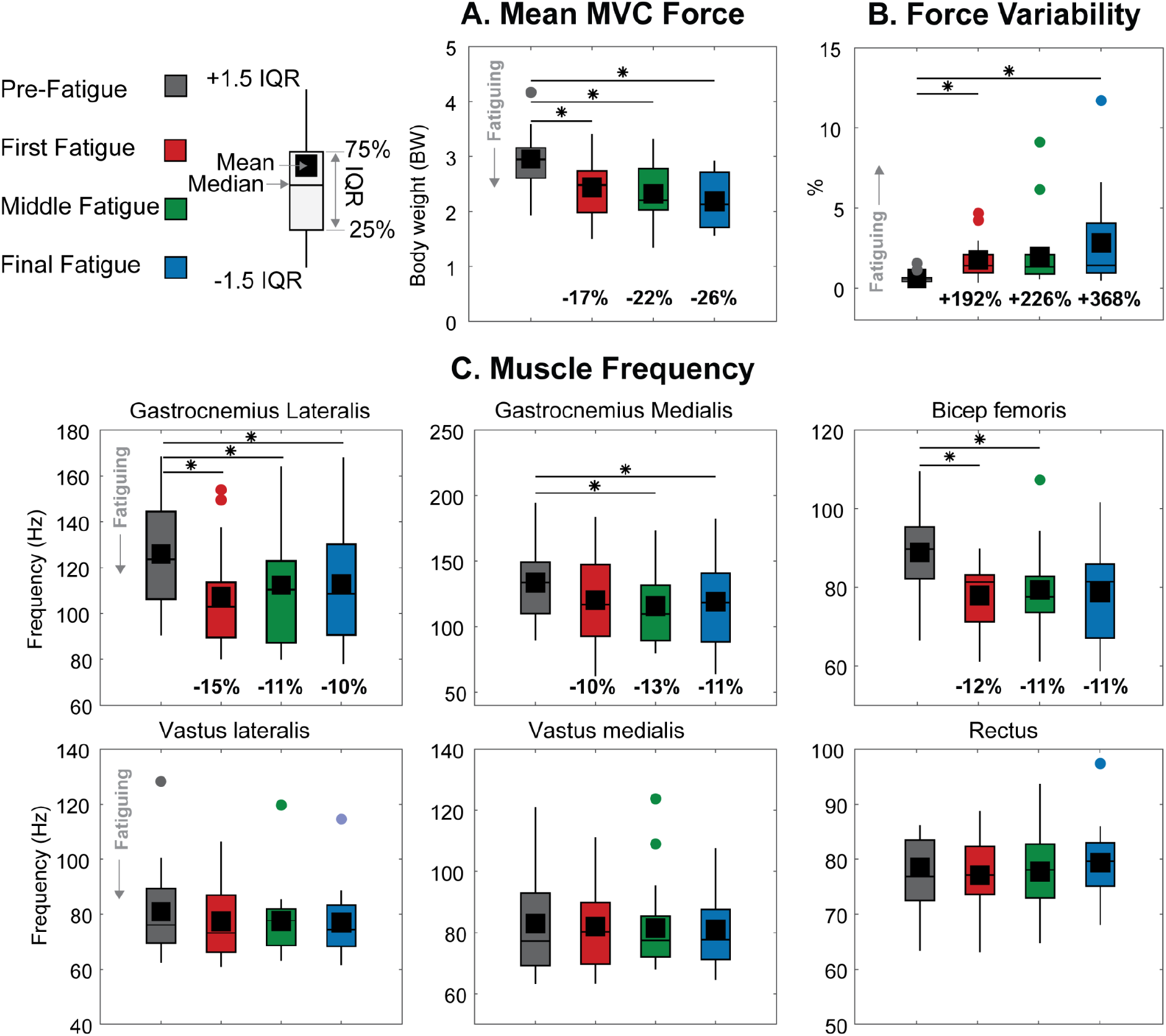
Objective measures of fatigue. As participants progressed through the protocol (Fatigue 1-3), they demonstrated A. A decline in Mean MVC vertical force, B. An increase in force variability, and C. A reduction in the median EMG frequency of several of the measured leg muscles. If we found the repeated measure ANOVA to be significant, we show any significant pairwise comparisons indicated with an asterisk. The legend on the top left explains the values shown in the box and whiskers plots.

### Fatigue led to reductions in leg force control responsiveness

As participants grew fatigued, their responsiveness decreased, demonstrated by a significant decline in both rise time (p = 0.0011) and bandwidth (p = 0.0014). However, this decrease in responsiveness wasn’t reflected in changes in fall time (p = 0.109) (Figure 6-A). Comparing the rise time of 303 ± 100 ms in the Pre-Fatigue condition to 373 ± 94 ms in the final fatigue condition, we observed a significant (p = 0.0006) increase of 23%. Simultaneously, bandwidth decreased significantly (p = 0.04831) from 1.33 ± 0.6 Hz to 1.00 ± 0.25 Hz, a reduction of 25%. Despite these changes, we did not observe any alteration in force control accuracy between these two conditions (Figure 6-B).

**Figure 6:**
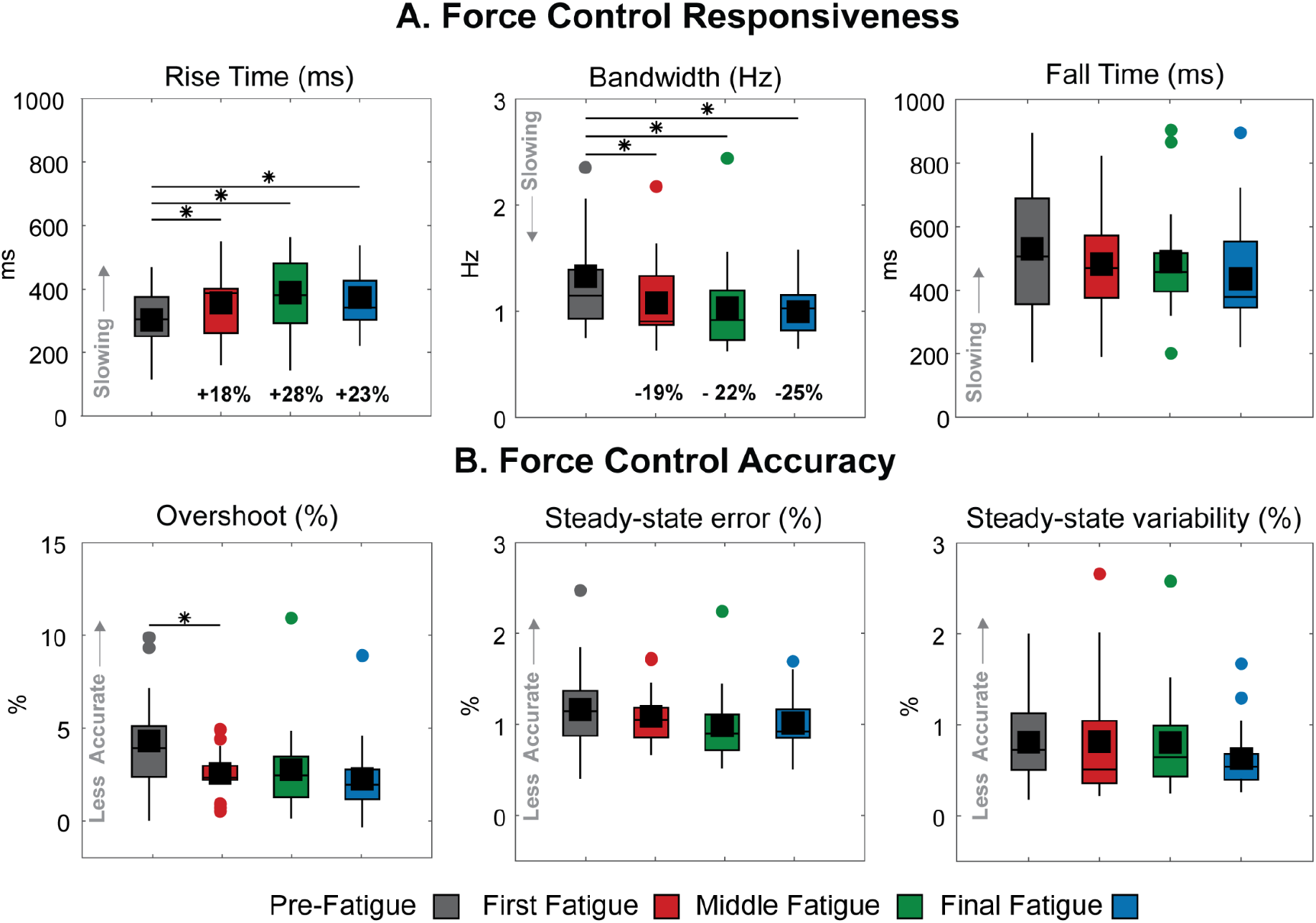
A. We assessed force control responsiveness by examining changes in rise time, bandwidth, and fall time. B. We evaluated force control accuracy through alterations in overshoot, steady-state error, and variability. In cases where we found the repeated measure ANOVA significant, we displayed significant pairwise comparisons, denoted by an asterisk.

### The Pulse Task was resilient to the effects of fatigue

As participants grew fatigued, they did not exhibit changes in responsiveness during the Pulse Task (Figure 7), evidenced by the absence of significant changes in rise time (p=0.9722), bandwidth (p=0.9147), or fall time (p=0.7830). Since the primary objective of the Pulse Task was responsiveness, we did not evaluate accuracy.

**Figure 7:**
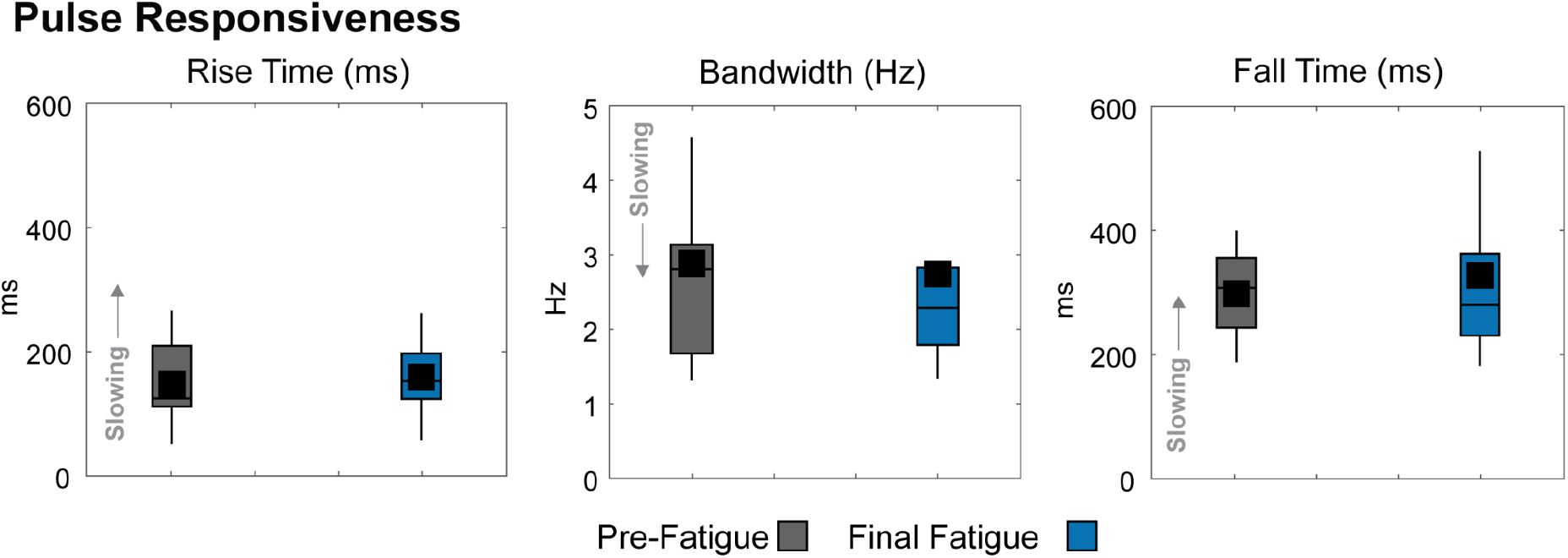
Pulse Task responsiveness, assessed by rise time, bandwidth, and fall time, comparing the Pre-Fatigue condition to the final fatigue condition, Fatigue-3. We did not evaluate the accuracy of the Pulse Task, as its primary objective was responsiveness. When the need for accuracy was removed from the objective, participants exhibited greater responsiveness compared to the Step Task, which required both responsiveness and accuracy.

### Participants exhibited higher responsiveness in the Pulse Task compared to the Step Task

We observed that participants demonstrated greater responsiveness in the Pulse Task than in the Step Task across all measures. In comparing the Pre-fatigue responsiveness of the two tasks, participants in the Pulse Task had a rise time of 148 ± 64 ms, a bandwidth of 2.88 ± 1.47 Hz, and a fall time of 299 ± 68 ms. In contrast, participants in the Step Task had a rise time of 303 ± 100 ms, a bandwidth of 1.33 ± 0.60 Hz, and a fall time of 531 ± 199 ms. During Pre-Fatigue, the Pulse Task exhibited a 51% faster rise time (p = 6.8e-6), a 116% wider bandwidth (p = 4.0e-4), and a 42% faster fall time (p = 6.0e-4) than the Step Task.

## Discussion

In this study, we investigated the impact of fatigue on the ability to control the vertical component of leg external ground forces, finding that fatigue led to a decrease in force control responsiveness (speed). Our protocol successfully fatigued participants, as shown by several objective measures including a decrease in mean maximum voluntary contraction force during the MVC Task, increased force variability during the Hold Tasks, and reductions in median EMG frequency in multiple lower leg muscles. Consistent with our hypothesis, we observed that fatigued participants demonstrated a significant decline in responsiveness, marked by a 23% increase in rise time and a corresponding 25% reduction in bandwidth. However, contrary to our hypothesis, we found that fatigued participants did not exhibit significant changes in leg force control accuracy. Furthermore, we found that in the Pulse Task—a task in which participants displayed overall greater responsiveness when compared to the Step Task—control was resilient to the effects of fatigue.

There are several limitations to our study. One limitation is that mechanical compliance, both within the participants’ bodies and within the testing apparatus, may have influenced the measured force. To minimize this potential issue, we used stiff materials and securely fixed the participants in the apparatus, as described in [1]. Another limitation is that we only evaluated force control at a single sub-maximal vertical force level. In real-life situations, leg external forces vary widely depending on the task. However, previous research suggests that aspects of force control might not differ significantly at different sub-maximal force levels of up to 60% of MVC when fatigued [1], [33]. A third limitation is that our study results may not generalize to other populations, such as older adults or individuals with chronic conditions. These groups could exhibit different control characteristics even before the onset of fatigue [34]. [35], [36]. Lastly, our study used a standardized fatigue protocol that might not accurately reflect real-world conditions in which fatigue occurs. For example, real-world conditions can involve not only physical but also cognitive demands, both of which may affect motor control and be subject to fatigue [37]. Our study targeted fatigue at the neuromuscular level and did not actively investigate the separate effects of physical and cognitive fatigue [38].

Fatigue can impact responsiveness through several pathways. One significant mechanism involves the inhibition of feedforward connections, or neural drive, from the brain to the muscles. Neural drive refers to the intensity or strength of the feedforward signals sent from the central nervous system to the muscles [39]. This reduction in neural drive may be due to a decrease in the release of neurotransmitters, an alteration in the firing patterns of motor neurons, or a reduction in the sensitivity of muscle fibres to neurotransmitters [39]. When an individual is fatigued, the intensity of these feedforward signals may be reduced, leading to a decrease in muscle force-generating capabilities [40], [41]. It may also be the case that fatigue leads to a reduction in the sensory feedback gains involved in the control of muscle force. In a feedback control system, responsiveness is determined, in part, by the ratio between output (e.g., muscle force) and sensory input (e.g., muscle length or velocity), known as feedback gains. Sensory feedback gains can be adjusted dynamically to achieve task success [42]. As feedback gains decrease, the control system becomes less responsive [43]. A comparison of the responsiveness results from the Step Task— which relies on feedback control and was affected by fatigue—and the Pulse Task—which relies more on feedforward control and was not affected by fatigue—suggests that in our experiment, fatigue primarily affected sensory feedback pathways and not neural drive.

The distinction between the effects of fatigue on feedback and feedforward control is important for understanding how the nervous system adapts to fatigue and could inform the development of interventions to maintain or improve motor performance under fatigued conditions [44]. For example, targeted interventions could focus on enhancing feedforward control through training programs [45] or provide external sensory feedback to optimize performance in the presence of fatigue [44]. The ability of individuals to maintain optimal performance in fatigued conditions is critical, particularly for athletes. Therefore, identifying and developing effective interventions can have a significant impact on their ability to perform at their best, even when fatigued.

Our observed reductions in leg force control responsiveness are large enough that they could negatively impact agile performance. Agile performance relies on the ability to quickly and accurately respond to changes in the environment. A decrease in responsiveness can increase the time needed for the system to respond accurately to these changes, leading to a decline in performance. This has been illustrated in simulation experiments using robotic legs, where reductions in responsiveness, as indicated by increases in rise time and narrowing of bandwidth, were observed when the controller time delay was increased [46]. And in hardware experiments with legged robots— narrowing bandwidth and increasing controller time delay resulted in the robot failing to land properly during a drop landing task [46]. However, the consequences of a 25% reduction in responsiveness, as observed in our study, on a system’s or individual’s agility may depend on their specific abilities and the task. For instance, the MIT Cheetah, a highly agile-legged robot, has a leg force responsiveness that is 100 times faster than that measured in humans—a 25% reduction in responsiveness may not significantly affect its performance [47], [48]. On the other hand, humans already exhibit comparatively slow rise times and narrow bandwidths, suggesting that a 25% decrease in responsiveness is more likely to impair agile performance. Indeed, fatigue-induced temporal changes in humans affect agile performance— fatigued athletes exhibit prolonged drop landing contact times [20] and decreased running speed [49].

Fatigue leads to reductions in leg force control responsiveness, potentially raising the risk of injury during physical activities. To preserve performance, individuals may compensate for these decreases by altering their body mechanics. For example, fatigued athletes have been observed landing from jumps with more flexed knees or shifting their load from plantar flexors to knee extensors [5]. Such fatigue-induced changes in body mechanics can compromise stability and heighten injury risk by placing extra stress on specific joints or muscles not typically engaged or active during certain movements [50], [51]. In men’s collegiate soccer matches, for example, players sustained approximately 50% more injuries in the second half compared to the first half [52].

In our experiment we developed a method for fatiguing participants and benchmarking the impact of fatigue on leg force control performance, laying groundwork for studying interventions that address fatigue and enhance agility. Potential interventions, such as strength training or exoskeletons, could be explored in diverse populations, including athletes, military personnel, and older adults. By understanding the effects of fatigue on leg force control responsiveness and accuracy, it may be possible to develop targeted strategies that not only mitigate the consequences of fatigue but also optimize agility across various tasks and populations. This research will help with future work focused on identifying, assessing, and implementing effective interventions that cater to the unique needs of individuals relying on leg force control for optimal performance.

## Acknowledgements

We would like to thank all the participants who volunteered their time for our study.

## Author Contributions

P.K. and J.M.D. conceived and designed research; P.K. performed experiments; P.K. analyzed data; P.K. J.M.W., S.N.R., and J.M.D. interpreted results of experiments; P.K. prepared figures; P. K. and J.M.D. drafted the manuscript; P.K., J.M.W., S.N.R., and J.M.D. edited and revised manuscript; P.K., J.M.W., S.NR, and J.M.D. approved the final version of the manuscript.

